# Spontaneous object exploration in a recessive gene knockout model of Parkinson’s disease: Development and progression of object recognition memory deficits in male *Pink1*-/- rats

**DOI:** 10.1101/2022.05.23.493123

**Authors:** Claudia C. Pinizzotto, Katherine M. Dreyer, Oluwagbohunmi A. Aje, Ryan M. Caffrey, Keertana Madhira, Mary F Kritzer

**Author notes:** **CORRESPONDENCE** Claudia C. Pinizzotto.

## Abstract

Cognitive impairments appear at or before motor signs in about one third of patients with Parkinson’s disease (PD) and have a cumulative prevalence of roughly 80% overall. These deficits exact an unrelenting toll on patients’ quality and activities of daily life due in part to a lack of available treatments to ameliorate them. This study used three well-validated novel object recognition-based paradigms to explore the suitability of rats with knockout of the PTEN-induced putative kinase1 gene (*Pink1*) for investigating factors that induce cognitive decline in PD and for testing new ways to mitigate them. Longitudinal testing of rats from three to nine months of age revealed significant impairments in male *Pink1*-/- rats compared to wild type controls in Novel Object Recognition, Novel Object Location and Object-in-Place tasks. Task-specific differences in the progression of object discrimination/memory deficits across age were also seen. Finally, testing using an elevated plus maze, a tapered balance beam and a grip strength gauge showed that in all cases recognition memory deficits preceded potentially confounding impacts of gene knockout on affect or motor function. Taken together, these findings suggest that knockout of the *Pink1* gene negatively impacts the brain circuits and/or neurochemical systems that support performance in object recognition tasks. Further investigations using *Pink1*-/-rats and object recognition memory tasks should provide new insights into the neural underpinnings of the visual recognition memory and visuospatial information processing deficits that are often seen in PD patients and accelerate the pace of discovery of better ways to treat them.

## INTRODUCTION

Parkinson’s disease (PD) is a common neurodegenerative disorder that is characterized by motor deficits such as bradykinesia, postural instability and resting tremor (Bloem, Okun et al. 2021, Vazquez-Velez and Zoghbi 2021). However, many PD patients also experience non-motor symptoms including impairments in cognition and memory (Aarsland, Bronnick et al. 2010, Aarsland, Creese et al. 2017, Goldman, Vernaleo et al. 2018, Fang, Lv et al. 2020). These impairments appear at or before motor signs in about one third of all PD patients and have a cumulative prevalence of more than 80% overall (Aarsland, Creese et al. 2017, Papagno and Trojano 2018, Fang, Lv et al. 2020). Although termed ‘mild cognitive impairments’ to distinguish these earlier occurring deficits from those associated with Parkinson’s disease-related dementia (PDD), these cognitive deficits exact a significant toll on patients’ quality and activities of daily life (Leroi, McDonald et al. 2012, Kudlicka, Clare et al. 2014, Oosterveld, Allen et al. 2015, Rodriguez-Blazquez, Forjaz et al. 2015, Barone, Erro et al. 2017, Saredakis, Collins-Praino et al. 2019). They also predict a more rapid and more severe clinical course of cognitive and motor decline and are associated with increased risk for freezing, falls and for developing PDD (Pigott, Rick et al. 2015, Mack and Marsh 2017, Cholerton, Johnson et al. 2018, Goldman, Vernaleo et al. 2018). Equally concerning is the lack of available treatments that can effectively combat these impairments (Goldman and Weintraub 2015, Mack and Marsh 2017, Goldman, Vernaleo et al. 2018, Fang, Lv et al. 2020). Using a series of novel object recognition-based paradigms, the studies presented here provide behavioral evidence indicating the potential suitability of rats bearing knockout of the PTEN (phosphatase and tensin homologue)-induced putative kinase1 gene (*Pink1-/-*) for facilitating the preclinical studies that are necessary to better understand and better treat cognitive and memory dysfunction in PD.

Currently, there are few available therapeutic options that are effective in preventing or slowing the progression of cognitive and memory decline in PD (Goldman and Weintraub 2015, Mack and Marsh 2017, Goldman, Vernaleo et al. 2018, Fang, Lv et al. 2020). Thus, in addition to clinical trials, animal and especially rodent models are being used to support controlled investigations into the risk factors and pathophysiological mechanisms that contribute to cognitive disturbance in PD and to test new ways of mitigating them (Fan, Han et al. 2021). There are several reasons to predict that *Pink1* -/- rats may be well-suited for these purposes. First, *Pink1*-/- rats have construct validity for the recessively inherited loss of function *Pink1* mutations that are the second most common mutation among autosomal recessive forms of PD; these mutations are also causally linked to early onset familial cases of PD (Valente, Abou-Sleiman et al. 2004, Kumazawa, Tomiyama et al. 2008, Scarffe, Stevens et al. 2014). In addition to disrupting mitochondrial function (Borsche, Konig et al. 2020), *Pink1* mutations in PD patients have also been shown to increase central nervous system vulnerability to reactive oxygen species, to dysregulate dopamine (DA) synthesis and reuptake (Gautier, Kitada et al. 2008, Bus, Zizmare et al. 2020, Goncalves and Morais 2021), to induce ferritin accumulation and iron toxicity in midbrain DA neurons (Hagenah, Becker et al. 2008) and to promote alpha-synuclein aggregation (Samaranch, Lorenzo-Betancor et al. 2010, Takanashi, Li et al. 2016, Nybo, Gustavsson et al. 2020). A rapidly growing literature documents characteristics similar to these in *Pink1* knockout rat lines (Urrutia, Mena et al. 2014, Villeneuve, Purnell et al. 2016, Creed and Goldberg 2018, Ren and Butterfield 2021). Further, although the data are not entirely consistent (de Haas, Heltzel et al. 2019), this rat strain has also been shown to undergo progressive loss of midbrain DA and brainstem norepinephrine (NE) neurons (Dave, De Silva et al. 2014, Grant, Kelm-Nelson et al. 2015, Villeneuve, Purnell et al. 2016, Cullen, Grant et al. 2018, Kelm-Nelson, Trevino et al. 2018). Earlier occurring, presumed compensatory changes in neostriatal concentrations and/or basal and potassium stimulated release of DA, acetylcholine (ACh), serotonin and other PD-relevant neurotransmitter systems have also been reported (Dave, De Silva et al. 2014, Creed, Menalled et al. 2019). Finally, *Pink1*-/- rats display behavioral deficits in motor and non-motor functions that mimic those experienced by PD patients. For example, in addition to age-related decline in gait coordination and grip strength(Dave, De Silva et al. 2014), *Pink1*-/- rats also demonstrate early-appearing deficits in sensorimotor cranial/otolaryngeal functions that negatively impact vocalizations, chewing and swallowing (Grant, Kelm-Nelson et al. 2015, Cullen, Grant et al. 2018, Kelm-Nelson, Brauer et al. 2018, Kelm-Nelson, Lechner et al. 2021). In addition, *Pink1* -/- rats also show behavioral correlates reflecting increased anxiety, e.g., changes in distress vocalizations, social approach, open vs. closed arm entries in elevated plus maze testing (Kelm-Nelson, Brauer et al. 2018, Cai, Qiao et al. 2019, Hoffmeister, Kelm-Nelson et al. 2021, Hoffmeister, Kelm-Nelson et al. 2022). This suggests face validity for the mood disturbances that are common in PD patients--including those with causal mutations in the *Pink1* gene (Ephraty, Porat et al. 2007, Ricciardi, Petrucci et al. 2014). However, there has been little systematic effort to determine whether *Pink1*-/- rats also model PD-relevant cognitive or memory phenotypes. This is despite evidence that among genetically determined forms of PD, patients with *Pink1* mutations have the greatest incidence of cognitive dysfunction and decline (Piredda, Desmarais et al. 2020, Gonzalez-Latapi, Bayram et al. 2021). To fill this gap in knowledge, longitudinal testing using Novel Object Recognition (NOR), Novel Object Location (NOL) and Object in Place (OiP) paradigms was used to determine whether and when *Pink1* -/- rats express deficits similar to the impairments in visual recognition memory and//or visuospatial information processing that commonly occur in PD patients (Owen, Beksinska et al. 1993, Higginson, Wheelock et al. 2005, Possin, Filoteo et al. 2008, Fang, Lv et al. 2020).

Object recognition-based behavioral paradigms are well-validated and widely used for evaluation of mnemonic constructs similar to those that are frequently at risk in neuropsychiatric disorders including Alzheimer’s disease, schizophrenia, PD and others (Grayson, Leger et al. 2015). Further, these single-trial tasks require no formal training, leverage spontaneous behaviors, are minimally stressful and can require minimal physical exertion (Ennaceur 2010, Luine 2015, Aggleton and Nelson 2020, Chao, de Souza Silva et al. 2020). These features are especially important for studying cognition and memory in preclinical models of PD where potentially confounding disease-related features of anhedonia, mood disturbance and motor impairment may be present. Finally, there is a rich, task-specific literature for object recognition paradigms describing the brain regions, networks and neurochemical systems that provide essential support for the different forms of recognition memories that these paradigms measure (Dere, Huston et al. 2007, Brown, Barker et al. 2012, Aggleton and Nelson 2020, Barker and Warburton 2020, Barker and Warburton 2020, Chao, de Souza Silva et al. 2020). Thus, there is a powerful interpretive framework at hand for gaining insights into the neural circuits that may be most affected by pathophysiology and where targeted therapeutics may be most beneficial. Given these benefits, it is not surprising that NOR, NOL and to a lesser extent OiP tasks continue to be widely used to assess cognitive deficits in a range of rodent models of PD (Grayson, Leger et al. 2015, Johnson and Bobrovskaya 2015, Haghparast, Esmaeili-Mahani et al. 2018, Ikram and Haleem 2019, Kyser, Dourson et al. 2019, Bharatiya, Bratzu et al. 2020, Boi, Pisanu et al. 2020, Fan, Han et al. 2021, Kakoty, K et al. 2021, Pinizotto 2022). This study extends this utilization for the first time to *Pink1*-/- rats in longitudinal comparative evaluations of object recognition memories in single cohorts of knockout and wildtype (WT) male Long Evans rats. In addition to object exploration and discrimination, the analyses presented below include assessments of motor function and affect that were made in conjunction with object recognition testing. Rats were also further evaluated at the beginning and end of the object recognition testing sequence using elevated plus maze testing, analyses of forelimb and hindlimb grip strength and assessments of foot slips in traversing a tapered balance beam.

## MATERIALS AND METHODS

### Animal subjects

Male Long Evans rats that were either wild type (WT, n=8) or *Pink1* knockouts [*Pink1* -/-, (LE-Pink1^em1Sage-/-^) n = 16] were purchased at 6 or 7 weeks of age (Envigo, Madison, WI, USA). All rats were double housed by genotype for the duration of the study in standard translucent tub cages (Lab Products, Inc., Seaford, DE, USA) filled with ground corn cob bedding (Bed O’ Cobs, The Anderson Inc., Maumee, Ohio, USA). Rats were kept under a 12-h non-reversed light-dark cycle with food (Purina PMI Lab Diet: ProLab RMH 3000) and water available *ad libitum*. Enrichment objects (Nyla Bones, Nylabone, Neptune, NJ USA) were also present in each cage. During the intervals when rats were not being behaviorally tested, they spent roughly 1 hour per week in groups of 2-6 in a large, dimly lit 6 ft square enclosure that contained tunnels, platforms and other larger scale objects for them to interact with. All procedures involving animals were approved by the Institutional Animal Care and Use Committee at Stony Brook University and were performed in accordance with the U.S. Public Health Service Guide for Care and Use of Laboratory Animals to minimize their discomfort. Rats were weighed every month as part of a measure of continued good health and to confirm an expected phenotype of greater body mass in age-matched *Pink1*-/- compared to WT control rats (Fig 1).

**Figure1.**
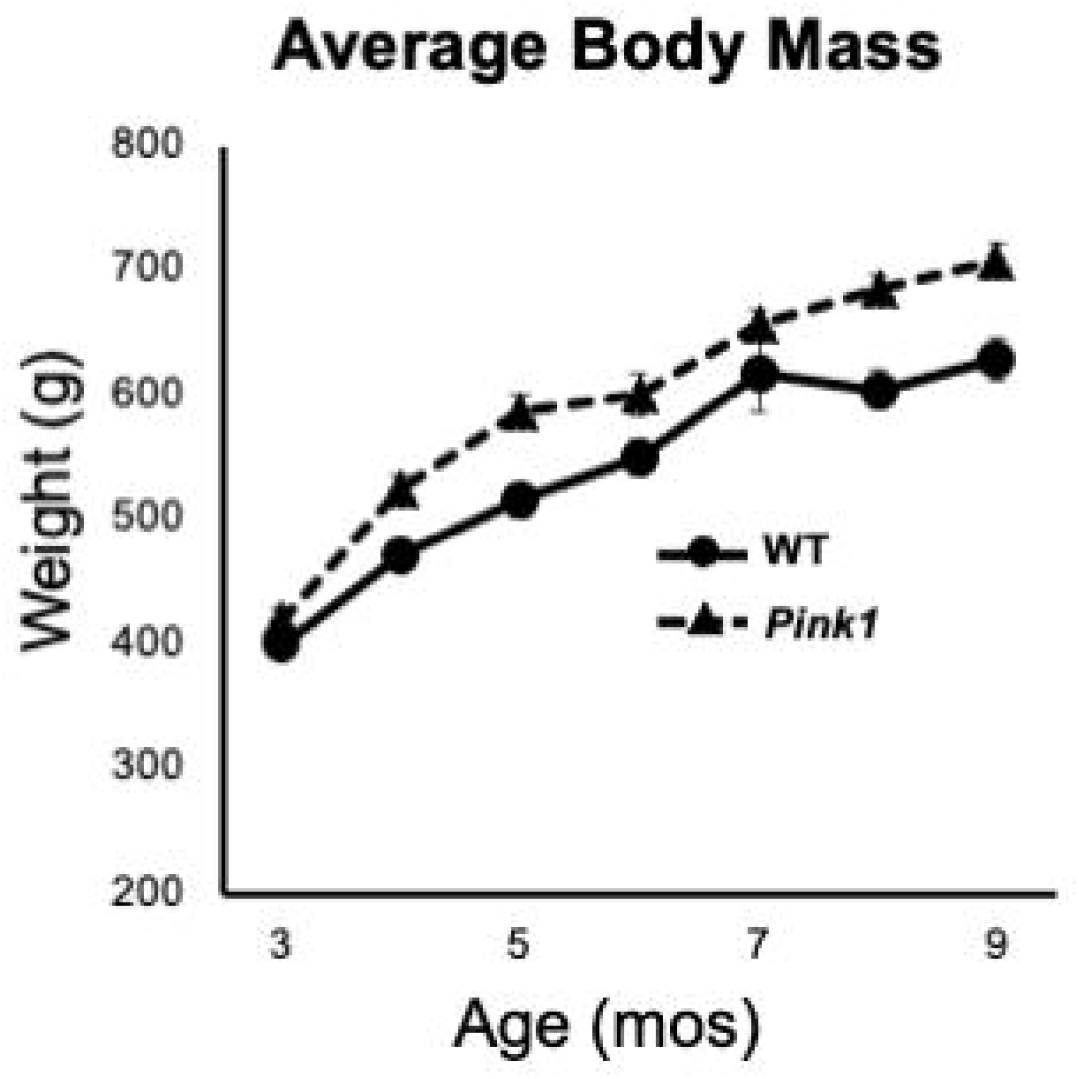
Line graphs showing changes in average weights in grams (g) of the male rats with knockout of the PTEN (phosphatase and tensin homologue)-induced putative kinase1 gene (*Pink1-/-*, triangles, dashed line) and the wild type (WT, circles, solid line) control male rats used in this study as they matured from 3 to 9 months (mos) of age. All rats continued to gain weight as the study progressed. As expected, the average weights of *Pink1*-/- rats were consistently greater than that of the WT cohort.

### Behavioral testing

Habituation and behavioral testing was conducted in a dedicated core facility that includes a central home cage holding room and 5 adjacent 10-12 ft square sound attenuated testing rooms. Each testing room had adjustable high contrast spatial cues on the walls and digital cameras to archive trials. Habituation and testing were conducted during rats’ subjective days between the hours of 9:00am and 1:00pm under ambient white lighting (∼ 260 lux).

### Apparatus

Object recognition tasks were carried out in open rectangular testing arenas (32 in long, 19 in wide,13 in high) made of translucent polypropylene. The arenas sat on a table 36in high. One of the long walls of the arena was made opaque and adjustable, small, high contrast cues were affixed to the outsides of the other three arena walls. These cues as well as distal room cues remained fixed during a given testing period and were rearranged across bimonthly testing sessions.

The elevated plus maze used was constructed of white laminate. It consisted of two open arms (5.5 in x 20.5 in), two closed arms (5.5 in x 20.5 in x 11.25) and an open central platform (5.5 in x 5.5 in). The maze was located 3 feet off the ground.

Grip strength was measured using a San Diego Instruments Animal Grip Strength System outfitted with two push/pull wire mesh force gauges (San Diego Instruments, San Diego, CA, USA).

The tapered balance beam used was composed of black plastic composite (Lafayette Instrument, Lafayette IN, USA). The top surface was rough to provide grip. The beam was 165 cm in length and tapered in width from 6 to 2 cm. Colored rulers were affixed to the sides of the beam that divided it into wide, medium and narrow thirds. A 2 cm wide ledge ran beside and below both sides of the beam surface to provide a crutch/step off position for rats to use if needed. Two digital cameras were used to record trials from the left and right sides that rendered all four feet visible.

### Habituation

One week after their arrival at Stony Brook University, rats were habituated to handling, to the central room of the testing facility and to gentle transfer between home cages and other enclosures. One week prior to the start of formal behavioral testing, rats were also habituated to the testing arenas and to the opaque start cylinders used in the object recognition tasks. This habituation consisted of daily exposures during which rats were placed in the start cylinder at the center of the arena; after 10 sec the cylinder was lifted and rats were given 10 min to explore the empty arena. This was repeated 2-3 times per day at roughly 60 min intervals for 5 days. The first round of object recognition testing began 3 days later; this and all subsequent rounds of object recognition testing began with an initial 5 min habituation trial in the empty arena.

### Testing procedures

Rats were behaviorally tested at 3, 5, 7 and 9 months of age on the NOR, NOL and OiP paradigms; these tasks were given pseudorandom order with 48 hours off in between each paradigm. Each trial began by placing rats in an opaque start cylinder located at the center of the arena. After a 10 sec delay, the cylinder was lifted and rats were free to explore. Different sets of sample and test objects were used for each task and for each time a given task was delivered. The arena and objects were cleaned with 70% EtOH before and after every trial.

#### Novel Object Recognition (NOR) Testing

Novel Object Recognition testing consisted of three 3 min sample trials, each separated by 1 hour, and one 3 min test trial separated from the last Sample Trial by 90 min. Rats were returned to home cages during intertrial intervals. During sample trials, two identical objects were placed in adjacent corners of the arena leaving at least 4-inch clearance from the walls. During the test trial rats explored objects that were in the same locations as during sample trials, albeit with one object from the sample trials and one that was novel. The objects that served as sample vs. novel objects and their locations in the arena were counterbalanced across rats in both groups.

#### Novel Object Location (NOL) Testing

Novel Object Location testing consisted of three 3 min sample trials each separated by 1 hour and a 3 min test trial separated from the last sample trial by 1 hr. Rats spent all intertrial intervals in home cages. During the sample trial, two identical objects were placed in adjacent corners of the arena with a 4-inch clearance from the walls. During test trials rats explored the same two objects but with one located in a corner that was occupied during the sample trial and the other placed in a previously unoccupied corner. The arena corners that served as sample vs. novel locations were counterbalanced across rats in both groups.

#### Object-in-Place (OiP) Testing

Object-in-Place testing consisted of three 3 min sample trials each separated by 5 min and a 3 min test trial separated from the last sample trial by a 5-minute inter-trial interval. Rats were returned to home cages during the intertrial intervals. For sample trials, 4 distinct objects were placed near each of the arena’s corners (4-inches from the walls). During Test trials, rats explored the same four objects, albeit with two occupying original positions and two occupying positions that were switched with each other. The positions and pairs of objects that occupied switched vs. stationary positions were counterbalanced across subjects in both groups.

#### Elevated Plus Maze testing

Rats were tested on the elevated plus maze at 3.5 and 9.5 months of age approximately 1 week after completing object recognition testing. At the start of the trial, rats were placed on the center portion of the maze facing away from the handler and were given a single 5 min trial to freely explore. All maze surfaces were cleaned with 70% ethanol before and after each trial.

#### Grip strength testing

Rats were held parallel to the center platform of the apparatus. Once they grasped the forelimb force plate, they were gently pulled backwards, away from it. After they released the forelimb plate, they continued to be drawn across the hindlimb force plate, which rats grabbed onto with hind feet while rats attempted to push forward. Thus, single trials were used to measure forelimb pull strength and hindlimb push/compressive strength. During each session, rats were given 3 trials that were separated by 30 sec to 1 min. The system automatically collects values of maximal force which were used for analyses.

#### Tapered Balance Beam testing

Wire tops were removed from rats’ home cages and the edge of the open home cage was used to support the narrow end of the balance beam. Rats were removed from the home cage and habituated to beam crossing by first placing them on the narrow end of the beam within a few steps of the home cage. Once they left the beam, they were given 1-2 min in the home cage as reward before being returned to the beam at wider and wider points (farther from the home cage). This was repeated until rats traversed the beam length with minimal stopping. All rats acquired this level of performance quickly, usually in less than 3 full length runs. Rats were then rested for about 15 min before being given 3 sequential full length trials.

### Data Analysis

Behavioral data were analyzed from digitally recorded trials by trained observers who were blind to genotype/group. Event-capture software (Behavioral Observation Research Interactive Software (BORIS) version 7.8.2, open access) was used to quantify the timing, instances and durations of specific behaviors defined below.

#### Object Recognition during Sample Trials

##### Total exploration

Total time in seconds rats spent actively exploring objects using vibrissae or snout. Sample trial object exploration was additionally evaluated for:

##### Spatial Bias

Total times in seconds rats spent actively exploring objects located in a given corner, quadrant or half of the arena.

##### Object Bias

Total times in seconds rats spent actively exploring distinct objects either presented simultaneously (OiP) or counterbalanced across subjects (NOR, NOL).

#### Object Recognition during Test Trials

##### Total exploration

Total time in seconds rats spent actively exploring objects using vibrissae or snout. Test trial object exploration was additionally evaluated for:

##### Discrimination index (DI)

###### NOR

Total time (in seconds) rats spent investigating novel (NO) vs familiar objects (FO), expressed as percent of total object exploration time. This index was calculated by the following formula:

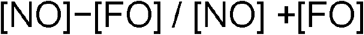

###### NOL

Total time (in seconds) rats spent investigating objects in new (Nw) vs original positions (Or), expressed as percent of total object exploration time. This index was calculated by the following formula:

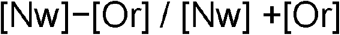

###### OiP

Total time (seconds) rats spent investigating two objects in switched (Sw) compared to original (Or) positions, expressed as percent of total object exploration time. This index was calculated by the following formula:

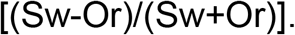

#### Other Behaviors

Test trials were analyzed for four major behaviors other than object exploration. The behaviors were defined as below:

- Rearing: total time (seconds) rats spent standing on hind paws either assisted by forepaw contact with objects or walls, or without assistance.
- *Grooming:* total time (seconds) rats spent preening any part of the head or body.
- Ambulation: total time (seconds) rats made forward motion via steps involving all four paws.
- Stationary: total time (seconds) rats sat at a given location and did not engage in grooming or object investigation.

#### Elevated Plus maze

Rats were evaluated for:

- Arm entries: Forward locomotion culminating in all four paws being inside a given arm. Separate counts were made of total arm entries, entries into closed arms and entries into open arms.
- Total times (in seconds) rats spent in open arms, closed arms or on the center platform of the maze.
- Duration (in percent total open arm occupation time) of major activities in open arms.
  - Head dipping- Investigation, with head and shoulders positioned over the edge of the open arm.
  - Ambulation- As per “Other Behaviors” above
- Duration (in percent total closed arm occupation time) of major activities in closed arms.
  - Rearing- As per “Other Behaviors” above
  - Grooming-As per “Other Behaviors” above
  - Ambulation- As per “Other Behaviors” above
  - Stationary-As per “Other Behaviors” above
- Duration (in percent total center platform occupation time) of major activities in center platform.
  - Stretch attend/scanning- total times rats spent making forward and back or side-to-side exploratory movements of the forebody with hindlimbs and tail remaining in place.
  - Rearing- As per “Other Behaviors” above
  - Ambulation- As per “Other Behaviors” above
  - Head dipping- Investigation with head and shoulders positioned over the edge of the open central platform.

#### Grip strength

The automated system was used to measure pull force of forelimbs and push/compressive force of hindlimbs. Values of maximal force recorded were normalized to body mass/weight prior to analysis.

#### Tapered balance beam

Rats were evaluated for foot slips made while traversing the full length of the balance beam. Data were collected separately for wide, middle and narrow portions of the beam. Because foot slips were rare for rats in both groups, these data were collapsed into measures of total numbers of foot slips per traversal for analysis. The percentage of rats per group committing some vs. no foot slips was also recorded.

### Statistics

Statistical analyses were performed using IBM SPSS, Version 25 (SPSS, Inc., Chicago, IL, USA). The data were first assessed for descriptive statistics, including Levine’s F-test for equality of variance. Comparisons of single measures across group/genotype were made using one-way analyses of variance (ANOVA), comparisons of measures across age were made using within-groups, one-way ANOVAs with repeated measures designs and comparisons of multiple measures made across groups used two-way repeated measures ANOVA. For all repeated measures comparisons, Mauchly’s test for sphericity of the covariance matrix was applied and degrees of freedom were adjusted as indicated using the Huynh-Feldt epsilon. Discrimination index data were additionally evaluated within groups using one sample t-tests to determine whether DI values were significantly different than zero, and relationships between individual measures of DI and rats’ total times spent exploring objects during sample and test trials were also assessed within groups by calculating Pearson’s correlation coefficients. All comparisons were additionally evaluated for effect sizes by calculating eta squared (*η*^2^) for ANOVAs or using Cohen’s D for t-tests.

## RESULTS

### Novel Object Recognition (NOR)

The pairs of objects used in NOR testing were similar in overall size and of relatively simple geometry; they differed along dimensions of color/contrast, composite material (plastic, glass or metal), surface features, e.g., smooth vs. grooved, and/or basic shape, e.g., cylindrical vs. rectangular.

#### Sample Trial Object Exploration

At 3 months of age, *Pink1*-/- rats spent nearly twice as much total sample trial time investigating objects as did WT controls (*Pink1*-/- = 93 sec, WT = 52 sec, Fig 2A); this difference was significant [ANOVA, F_(1,21)_ = 26.61, p< 0.001, □^2^ = 0.56]. During subsequent testing *Pink1*-/- rats also tended to spend more time exploring sample objects (Fig 2A). However, the differences in sample observation times seen at these later ages were relatively small (5 months = 142 vs. 128 sec; 7 months = 62 vs. 56 sec; 9 months = 75 vs. 75 sec) and were not significantly different across genotype (□^2^ = 0.01-0.05).

**Figure 2.**
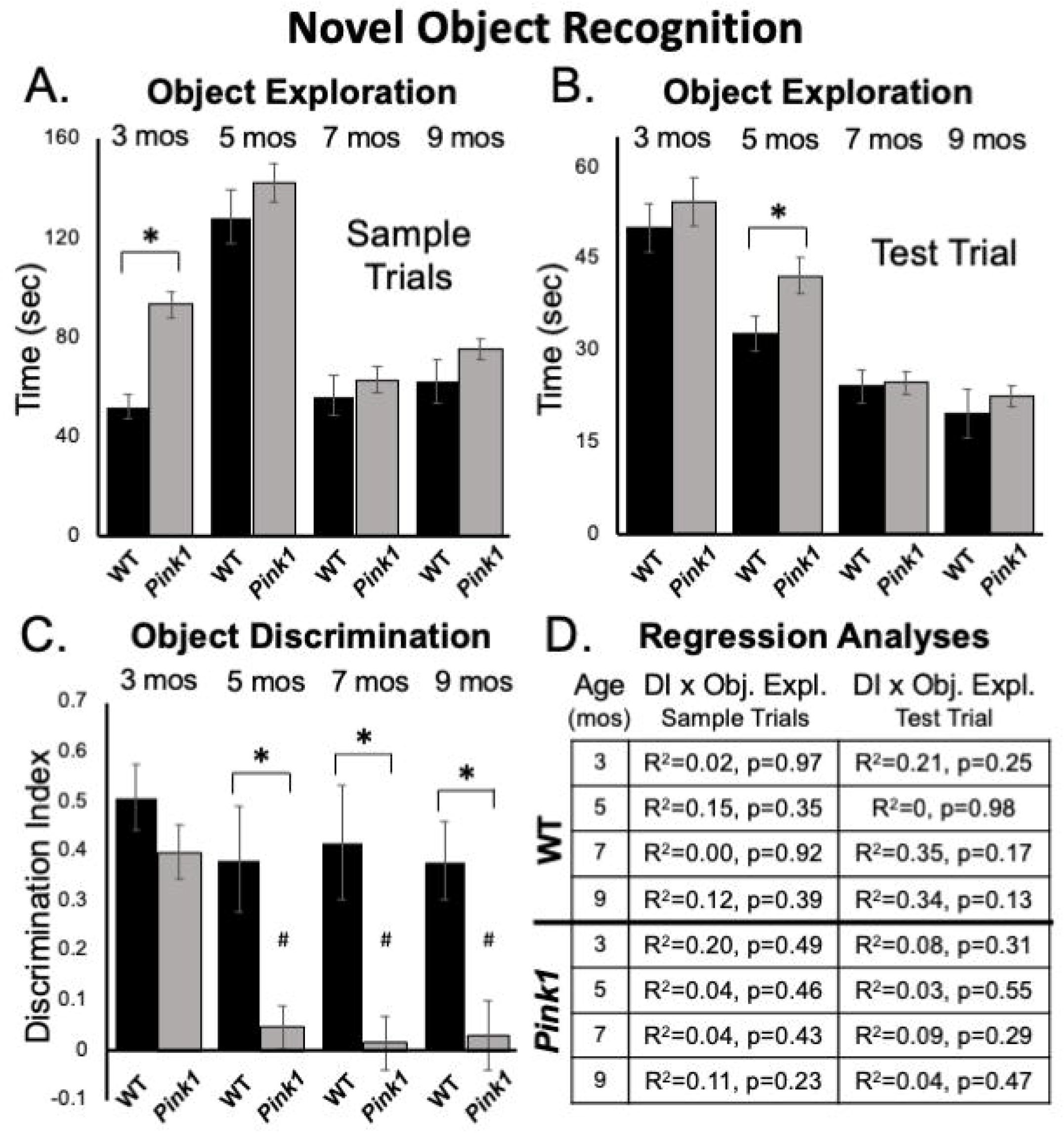
Novel Object Recognition data. Bar graphs showing total times in seconds (sec) that wild type (WT, black bars) and rats with knockout of the PTEN (phosphatase and tensin homologue)-induced putative kinase1 gene (*Pink1-/-*, gray bars) spent actively exploring objects during Sample Trials (A) and during Test Trials (B) in testing at 3, 5, 7 and 9 months (mos) of age. Overall, *Pink1*-/- rats spent more time than WT rats investigating objects; asterisks identify these differences as significant at 3 mos of age for Sample Trials (A), and for 5 mos of age for Test Trials (B). (C) Bar graphs showing calculated discrimination index (DI) for WT (black bars) and *Pink1*-/- rats (gray bars). This measure of object recognition memory was similar in rats of both genotypes at 3 months of age. At all other ages, DIs were significantly greater (*) in WT than in *Pink1*-/- rats. Within-groups comparisons of DI across age showed that in *Pink1*-/- rats, DIs measured at 5, 7 and 9 months of age were significantly lower (#) than DI measured at 3 months of age. (D) Tables showing R^2^ and p values for regression analyses that compared DIs to total sample object exploration times and to test trial object exploration times in WT and *Pink1*-/- rats at each age tested. No significant or near significant correlations were found among these measures.

#### Sample Trial Object Bias

At all ages tested, rats in both groups tended to divide total times investigating sample objects more or less equally among the two objects present. This was confirmed in a series of within-groups repeated measures ANOVAs that in most cases found no significant main effects of object position on object exploration (3 *Pink1*-/-, WT, □^2^ = 0.007, 0,013; 9 months: *Pink1*-/-, WT, □^2^ = 0.023, 0.15). The only exception occurred among WT controls at 5 months of age. For this timepoint, a significant main effect of Object Position (F_(1,2)_ = 9.16, p= 0.019, □^2^ = 0.57) was found that was driven by WT rats dividing total sample object investigation times among the two objects present according to a ratio of roughly 60 to 40. There were no significant group differences noted in sample trial object exploration times based on which object was used as sample at any age (□^2^ = 0-0.064).

#### Test Trial Object Exploration

Rats of both genotypes spent roughly 10-30% of test trial times exploring objects (Fig 2B). On average *Pink1*-/- rats spent more time exploring objects than WT controls (Fig 2B). However, these differences were generally less than 5 seconds (3 months = 54 vs. 50 sec; 5 months = 42 vs. 33 sec; 7 months = 25 vs. 24 sec; 9 months = 22 vs. 20 sec). Analyses of variance showed that main effects of genotype on this measure were only significant at 5 months of age [F _(1,21)_ = 4.44, p = 0.047 □^2^ = 0.17].

#### Test Trial Object Discrimination

Rats of both genotypes demonstrated robust discrimination of novel compared to familiar objects at 3 months of age (WT DI = 0.51; *Pink1*-/- DI = 0.40, Fig 2C). An ANOVA confirmed that there were no significant differences between these two values (□^2^ = 0.068) and one-sided t-tests showed that DI values for rats of both genotype were significantly different/greater than zero [WT: *t*(7) = 7.56, *Pink1*-/-: *t*(14) = 7.26, p < 0.001, *d* = 0.19 and 0.21, respectively]. During subsequent testing, WT rats maintained robust levels of novel object discrimination (Dis = 0.38 - 0.42, Fig 2C). A within-groups, repeated measures ANOVA further confirmed that DI values in this group were unchanged from 3-9 months (□^2^ = 0.11), and one-sided t-tests showed that all DI values were significantly different/greater than zero [*t*(7) = 3.6-4.8, p < 0.001-0.006, *d* = 0.22-0.31]. In contrast, novel object discrimination in 5-month-old *Pink1-/-* rats dropped dramatically to very low levels that were maintained up to 9 months of age (DIs = 0.02-0.05, Fig 2C). A within-groups ANOVA identified significant impacts of age on DI in the *Pink1-/-* group (F_(3,42)_ = 11.12, p<0.001, □^2^ = 0.44) and follow-up comparisons confirmed that the DIs measured at 5, 7, and 9 months in these knockout rats were significantly lower than that measured at 3 months (p<0.001 for all ages). A series of one-sided t-tests also showed that none of the DI values measured in *Pink1*-/- rats at 5-9 months of age were significantly different than zero (*d* = 0.17-0.27). Finally, ANOVAs that compared groups identified main effects of genotype on DI in rats that were significant at 5, 7 and 9 but not 3 months of age (5 months: F_(1,21)_ = 12.05, P = 0.002; 7 months F_(1,20)_ = 13.00, p = 0.002; 9 months: F_(1,21)_ = 9.65, p = 0.005, □^2^ = 0.32-0.39, Fig 2C). Regression analyses confirmed that there were no significant or near significant positive correlations between DIs and measures of object exploration during sample or test trials for either genotype at any age (R^2^ = 0-0.35, p = 0.13-0.98, Fig 2D).

#### Test Trial

*Other Behaviors*. Analyses of ambulation, rearing, grooming and remaining stationary during NOR test trials identified significant main effects in the way that animals apportioned test trial times across these behaviors [F _(1.52-2.20, 30.92-56.60)_ = 17.43-33.41, p < 0.001 for all, □^2^ = 0.45-0.61]. For all testing except at 7 months of age, significant main effects of genotype [F_(1, 21)_ = 4.66, p = 0.043, □^2^ = 0.18] and/or significant interactions between genotype and behavior [F_(1.194-2.69, 40.20-56.57)_ = 3.76-9.64, p <0.001-.027, □^2^ = 0.15-0.32) were also found. Follow up comparisons further showed that at every testing age *Pink1*-/- rats spent significantly less time grooming than the WT rats (3 months = 1.4 vs. 25 sec, p < 0.001; 5 months = 6.3 vs. 25 sec, p < 0.001; 9 months = 11 vs. 32 sec, p = 0.001). At 9 months of age *Pink1*-/- rats were also found to spend significantly more time ambulating (29 vs. 19 sec, p = 0.017) and rearing (57 vs. 29 sec, p = 0.002) and significantly less time remaining stationary (60 vs. 81 sec, p = 0.011) than WT controls. At all other testing ages, rats of both genotypes spent similar amounts of NOR test trial times engaged in these activities.

### Novel Object Location (NOL)

Because there is no need to match valence between objects within trials, the pairs of objects used in NOL had more complex geometric features, e.g., depressions, spurs, sharp corners, than objects used for the NOR or OiP paradigms.

#### Sample Trial Object Exploration

At 3 months of age, an ANOVA confirmed that the *Pink1*-/- rats spent significantly more total sample trial times investigating objects than WT subjects [121 vs. 82 sec, F_(1,21)_ = 7.05, p= 0.015, □^2^ = 0.25, Fig 3A]. However, group differences (*Pink1*-/- vs. WT) in sample object exploration at subsequent ages were all negligible (5 months = 54 vs. 55 sec; 7 months = 39 vs. 40 sec; 9 months = 48 vs. 53 sec) and were not significant (□^2^ = 0.001-0.013).

**Figure 3.**
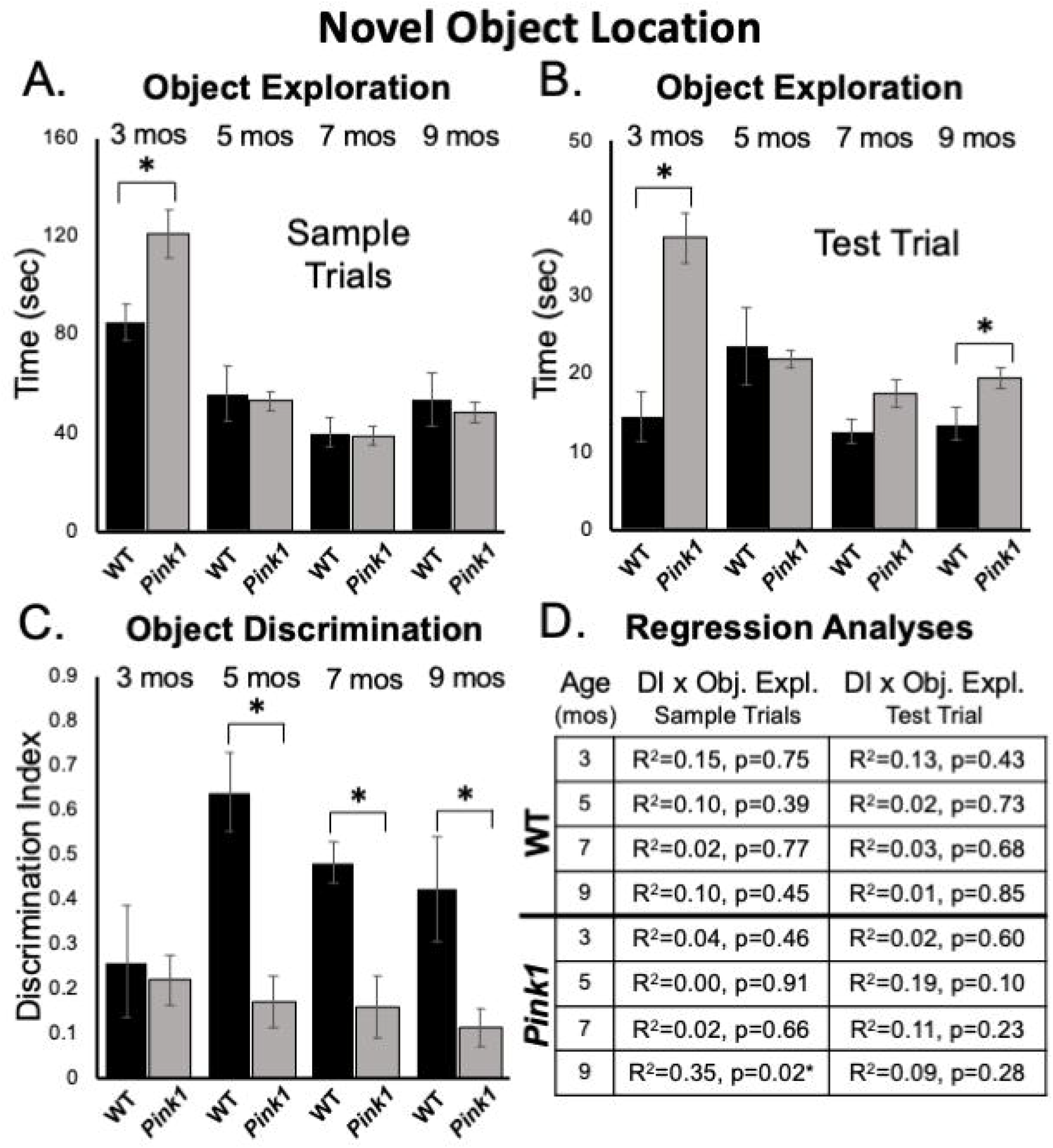
Novel Object Location data. Bar graphs showing total times in seconds (sec) that wild type (WT, black bars) and rats with knockout of the PTEN (phosphatase and tensin homologue)-induced putative kinase1 gene (*Pink1-/-*, gray bars) spent actively exploring objects during Sample Trials (A) and during Test Trials (B) in testing at 3, 5, 7 and 9 months (mos) of age. In general, *Pink1*-/- rats spent equal or more time than WT rats investigating objects; asterisks identify object exploration times as significantly greater in the *Pink1*-/- compared to WT cohort for testing at 3 mos of age during Sample Trials (A), and for testing at 3 and 9 mos of age during Test Trials (B). (C) Bar graphs showing calculated discrimination index (DI) for WT (black bars) and *Pink1*-/- rats (gray bars). This measure of object location memory was similar in rats of both genotypes at 3 months of age. At all other ages, DIs were significantly greater (*) in WT than in *Pink1*-/- rats. (D) Tables showing R^2^ and p values for regression analyses comparing DIs to total sample object exploration times and to test trial object exploration times in WT and *Pink1*-/- rats at each age tested. No significant or near significant positive correlations were found among these measures. However, a significant negative correlation (*) between increased sample trial object exploration and lower DI values was found for *Pink1*-/- rats at 9 months of age.

#### Sample Trial Object Bias

Analyses of total sample trial object explorations as functions of object position showed that rats of both genotypes investigated the two sample objects present to similar extents. The largest difference seen in exploring one vs. the other object was for 3-month-old WT rats, where an average difference on the order of about 10 sec was seen. However, within-groups repeated measures ANOVAs confirmed that this difference and most others were not significant (□^2^ = 0.001-0.22). The single exception was for 9 month old WT rats, where relatively small differences in the amounts of times spent investigating objects located in each the two corners (23 vs. 29 sec) proved significant [F(_1,7)_ = 5.96, p= 0.045, □^2^ = 0.46].

#### Test Trial Object Exploration

Analyses of total times spent exploring objects during NOL test trials showed that *Pink1*-/- rats generally spent more time investigating objects than the WT controls (3 months = 38 vs. 14 sec; 5 months = 21 vs. 24 sec; 7 months = 18 vs. 13 sec; 9 months = 19 vs. 13 sec, Fig 3B). These group differences were significant for rats at 3 [F _(1,21)_ = 19.58, p < 0.001, □^2^ = 0.50] and 9 months of age [F_(1,21)_ =6.06, p = 0.023, □^2^ = 0.22] but did not reach a critical difference at the other two testing ages (□^2^ = 0.01, 0.15).

#### Test Trial Object Discrimination

At 3 months of age, rats of both genotypes showed modest discrimination of objects in novel compared to familiar locations (WT DI = 0.26; *Pink1*-/- DI = 0.22, Fig 3C). Thereafter, object location discrimination rose to and remained at considerably higher levels in WT rats for the duration of testing (DI = 0.49-0.64, Fig 3C). In contrast, NOL discrimination in the *Pink1*-/- group remained low across all subsequent testing (DI = 0.12-0.17, Fig 3C). Nonetheless, one sample t-tests showed that all DI values for both groups were significantly different/higher than zero [WT: *t*(7) = 2.10- 10.54 p < 0.001- 0.04, d = 0.13- 0.33; *Pink1*-/-: *t*(14)= 2.37- 3.83, p <0.001- 0.017, d = 0.17- 0.25]. Within-groups, repeated measures ANOVAs also showed that there were no significant main effects of age on DIs for rats of either genotype (□^2^ = 0.059-0.25). However, across-groups ANOVAs confirmed that DIs in WT rats were significantly higher than those of the *Pink1*-/- cohort at 5, 7 and 9 months of age [F_(1,21)_ = 11.18-21.28, p < 0.001- 0.003, □^2^ = 0.36-0.50, Fig 3C]. Regression analyses also confirmed that for both groups there were no significant or near significant positive correlations between DIs and measures of sample or test trial object exploration at any age (R^2^ = 0.001-0.19, p = 0.10-0.91, Fig 3D). However, at 9 months of age a significant negative correlation (greater object exploration/lower DI) was identified between total sample trial objective exploration and DI in the Pink1-/- group [F_(1,13)_ = 7.11, p = 0.019, R^2^ = 0.35, Fig 3D].

#### Test Trial

*Other Behaviors*. Analyses of ambulation, rearing, stationary and grooming revealed significant differences in the amounts of time rats of all ages allotted to these activities [F _(1.46-2.15, 27.81-45.15)_ = 21.56-43.74, p < 0.001 for all, □^2^ = 0.53-0.68]. Significant main effects of genotype [F_(1, 20)_ = 5.40-13.13, p = 0.002-0.031, □^2^ = 0.22-0.40] and/or significant interactions between genotype/group and behavior [F_(1.55-2.15, 30.93-45.15)_ = 5.67-14.40, p <0.001-.005, □^2^ = 0.21-0.42] were also identified at all testing ages. Although there was some variance in the data, in general, main effects were driven by *Pink1*-/- rats spending more time engaged in active behaviors (rearing, ambulation) and less time being sedentary (stationary, grooming) than WT rats. This was borne out in follow up comparisons that showed *Pink1* -/- rats groomed significantly less than the controls at all ages (3 months = 4 vs. 26 sec, p < 0.001; 5 months = 5 vs. 17 sec, p = 0.002; 7 months = 3 vs. 22 sec, p= 0.001; 9 months = 11 vs. 32 sec, p < 0.001) and spent significantly less time stationary than WT rats in testing at 3 and 5 months of age (3 months = 33 vs. 61 sec, p< 0.001; 5 months = 32 vs. 64 sec, p = 0.004). The *Pink1 -/-* group also spent significantly more time ambulating than WT controls in testing at 5 and 7 months of age (5 months = 39 vs. 29 sec, p = 0.017; 7 months = 38 vs. 28 sec, p = 0.004) and significantly more time rearing at 5 months of age (81 vs. 44 sec, p = 0.001).

### Object in Place (OiP)

The groups of objects used in OiP testing were closely matched in terms of simple geometry and overall size but differed in properties that included color/contrast, material (plastic, glass or metal), general shape and/or subtle surface features.

#### Sample Trial Object Exploration

At 3 months of age, *Pink1-/-* rats spent significantly more time investigating sample objects than WT rats [127 vs. 91 sec, F_(1,21)_ =11.73, p= 0.003, □^2^ = 0.36, Fig 4A]. However, at 5 months of age, WT rats spent significantly more time investigating the samples than the *Pink1*-/- cohort [111 vs. 88 sec, F _(1, 21)_ = 7.66, p = 0.012, □^2^ = 0.27, Fig 4A]. In testing at 7 and 9 months of age there were no significant main effects of group/genotype on total sample object exploration times between *Pink1*-/- and WT rats (7 months = 74 vs. 85 sec, □^2^ = 0.09; 9 months = 77 vs. 80 sec, □^2^ = 0.005, Fig 4A).

**Figure 4.**
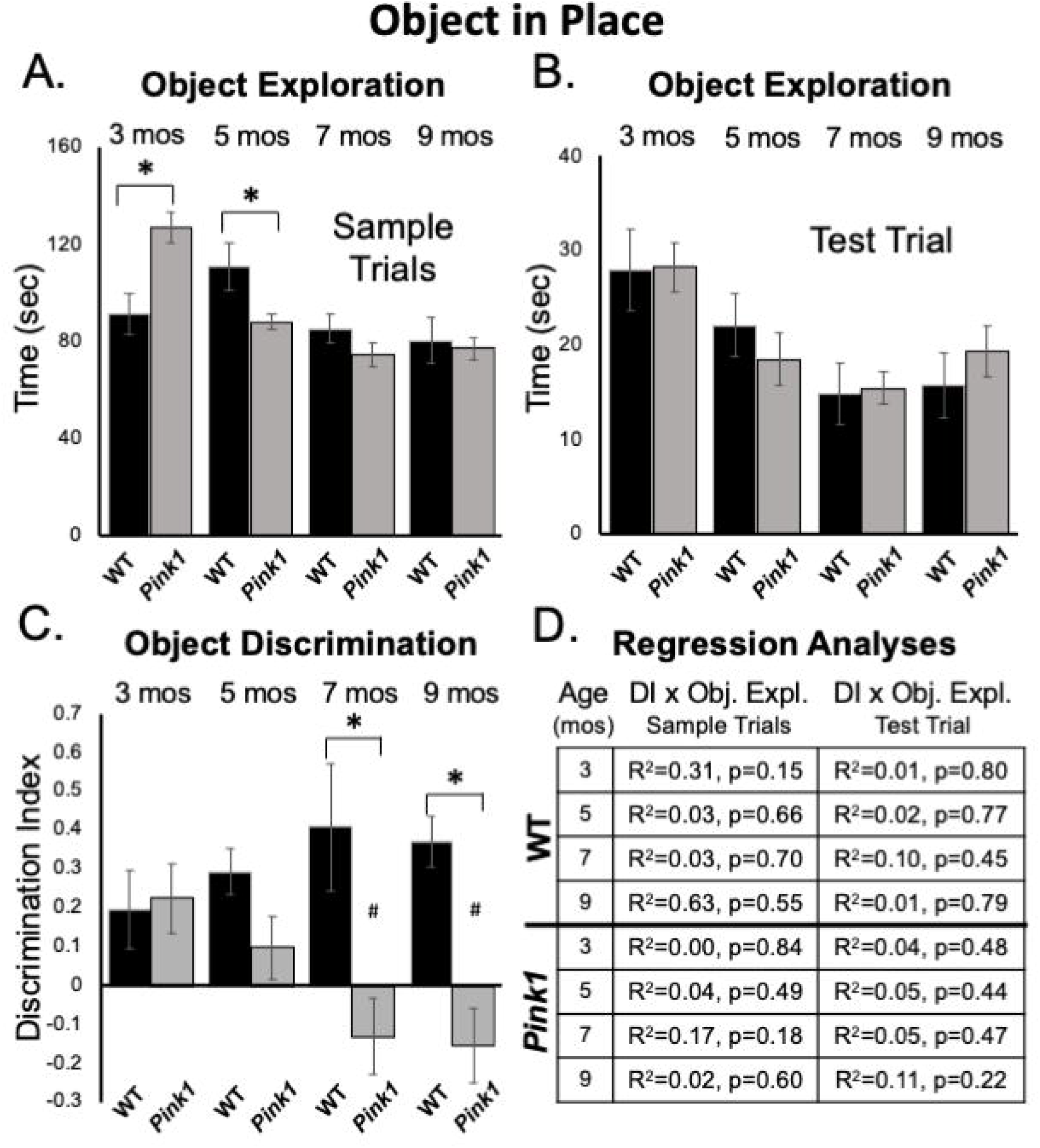
Object-in-Place data. Bar graphs showing total times in seconds (sec) that wild type (WT, black bars) and rats with knockout of the PTEN (phosphatase and tensin homologue)-induced putative kinase1 gene (*Pink1-/-*, gray bars) spent actively exploring objects during Sample Trials (A) and during Test Trials (B) in testing at 3, 5, 7 and 9 months (mos) of age. In general, the amounts of time spent exploring objects were comparable among the *Pink1*-/- WT groups. However, asterisks in (A) identify object exploration times that were significantly greater in the *Pink1*-/- compared to WT cohort for testing at 3 mos of age, and that were significantly greater in the WT compared to *Pink1*-/- rats for testing at 5 mos of age during. (C) Bar graphs showing calculated discrimination index (DI) for WT (black bars) and *Pink1*-/- rats (gray bars). This measure of integrated object recognition memory was similar in rats of both genotypes at 3 and 5 months of age. At all other ages, DIs were significantly greater (*) in WT than in *Pink1*-/- rats. Within groups comparisons of DI across age showed that in *Pink1*-/- rats, DIs measured at 7 and 9 months of age were significantly lower (#) then DI measured at 3 and 5 months of age. (D) Tables showing R^2^ and p values for regression analyses comparing DIs to total sample object exploration times and to test trial object exploration times in wild type and *Pink1*-/- rats at each age tested. No significant or near significant correlations were found among these measures.

#### Sample Trial Object Bias

Rats in both groups investigated each of the four sample items present approximately equally and divided observation times similarly across objects located in each of the arena’s four corners. Thus, there were no indications of bias based on object type or position. This was confirmed in a series of within-groups, repeated-measures ANOVAs that found no significant main effects of object type or arena corner *(Pink1*-/-: □^2^ = 0.011-0.11; Control: □^2^ = 0.008-0.24) on measures of object exploration.

#### Test Trial Object Exploration

Analyses of total times spent exploring objects during test trials showed that *Pink1*-/- and Control rats both spent similar amounts of test trial times exploring objects (3 months = 28 sec, both; 5 months = 19 vs. 22 sec; 7 months = 15 sec, both; 9 months = 19 vs. 16 sec, Fig 4B). There were no significant main effects of genotype on this measure (□^2^ = 0.00-0.031).

#### Test Trial Object Discrimination

At 3 months of age, WT and *Pink1*-/- rats both showed similar ability to discriminate among objects located in exchanged compared to original positions (WT DI =0.20; *Pink1*-/- DI = 0.22, Fig 4C); one sample t-tests showed that all DI’s measured in both groups were significantly different/greater than zero [WT: *t*(7) = 1.95, p = 0.046 and d = 0.28; *Pink1*-/-: *t*(13) = 2.52, p = 0.013 and d = 0.33, respectively]. However, testing at later time points showed that DIs in WT rats tended to incrementally increase (5 months DI = 0.29; 7 months DI =0.41; 9 months DI = 0.37, Fig 4C). One sample t-tests confirmed that all WT DI values were significantly different/greater than zero [*t*(7) = 2.49- 5.77, p < 0.001- 0.021, d = 0.18- 0.46]. However, within-groups repeated measures ANOVAs showed that the incremental increases in DI observed across age were not significant (□^2^ = 0.09). In contrast, average DIs in the *Pink1*-/- group showed a stepwise decline from 5 to 9 months of age (5 months DI = 0.10; 7 months DI =-0.13; 9 months DI = -0.15, Fig 4C). One sample t-tests showed that DI’s measured across this interval were not significantly different than zero (d = 0.32 - 0.37) and a within-groups repeated measures ANOVAs confirmed DI’s significantly declined with age [F _(3,30)_ = 5.18, p < 0.005, □^2^ = 0.34). Follow-up comparisons specifically identified DI’s measured at 3 and 5 months as significantly greater than those measured at 7 and 9 months of age (p =0.003-.047, Fig 4C). Finally, across-groups ANOVAs showed that while DIs between the WT and *Pink1*-/- rats were initially similar, their diverging trajectories culminated in significant group differences at 7 and 9 months of age [F_(1,21)_ = 8.94-13.95, p< 0.001-0.008, □^2^ = 0.33-0.40, Fig 4C). Importantly, regression analyses confirmed that there were no significant or near significant positive correlations between DI and measures of object exploration during sample or test trials for any group at any age (R^2^= 0.004-0.31, p= 0.15-0.84, Fig 4D).

#### Test Trial: Other Behaviors

Analyses of ambulation, rearing, stationary behavior and grooming revealed significant differences in the amounts of time rats of all ages allotted to these activities [F _(1.39-3, 23.59-60)_ = 12.56-90.36, p < 0.001 for all, □^2^ = 0.43-0.81]. However, there were no significant main effects of genotype (□^2^ = 0-0.047) and no significant interactions between genotype/group and behavior (□^2^ = 0.03-0.12) at any testing age.

### Elevated Plus Maze

At 3.5 months of age, rats in both groups made comparable numbers of total arm entries (WT = 9.14, *Pink1*-/- = 9.60) that for both genotypes were biased toward entries into closed vs. open arms by more than 2 to 1 (Fig 5A). However, a repeated measures ANOVA that compared times spent in a given sector of the maze (open arms, closed arms, center platform, Fig 5B) identified significant main effects of maze location [F_(2,40)_ = 89.12, p <0.001, *η*^2^ = 0.82] and a significant interaction between maze location and genotype/group [F_(2,40)_ = 11.99, p < 0.001, *η*^2^ = 0.38]. Follow-up pairwise comparisons showed that these effects were driven by WT rats spending significantly more time in closed arms (129 vs. 85 sec, p = 0.005) and significantly less time in the maze center (136 vs. 190 sec, p < 0.001) compared to the *Pink1*-/- cohort (Fig 5B). Analyses of major behaviors exhibited within each maze location (expressed as percent total times spent within these sectors) also revealed group differences (Fig 5C-E). For all three maze compartments, significant main effects were identified for times allotted to particular major behaviors [F_(1.61-2.34, 32.10-46.69)_ = 42.95-51.48, p < 0.001 for all, *η*^2^ = 0.68-0.72]. However, significant interactions between group and compartment specific behaviors were only significant for open [F(1,20) = 18.73, p < 0.001, *η*^2^ = 0.48, Fig 5C] and closed [F(2.34,46.69) = 9.18, p <0.001, *η*^2^ = 0.32, Fig 5D] arm locations. For the open arms, main effects were driven by WT rats spending about 30% more time head-dipping than *Pink1*-/- rats (p<0.001, Fig 5C). For the closed arms, these effects were driven by WT rats spending roughly 12% less time rearing and 16% more time grooming compared to *Pink1*-/- rats (p< 0.001 for both, Fig 5D). In the center platform, WT and *Pink 1*-/- rats engaged in stretch attend/scanning (41.6, 49.3% of time), ambulating (14.6, 13.4% of time), head dipping (15.7, 9.5% of time) and rearing (10.0, 13.7% of time) similarly (Fig 5E). There were no significant group differences in these allotted times (*η*^2^ = 0.065, 0.001)

**Figure 5.**
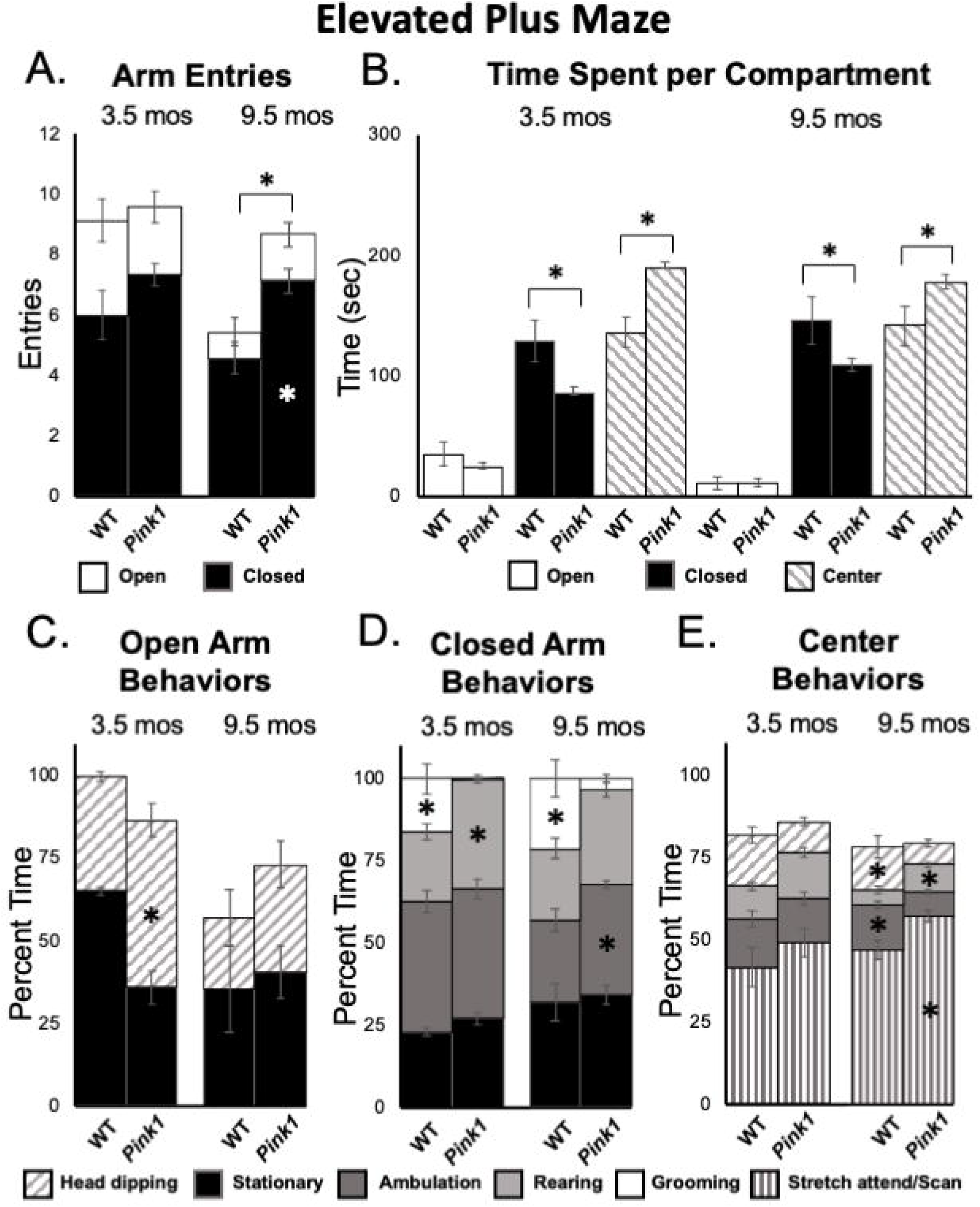
Elevated Plus Maze data. (A) Stacked bar graphs showing total arm entries divided into total entries made into open (white) and closed (black) arms for wild type (WT) and rats with knockout of the PTEN (phosphatase and tensin homologue)-induced putative kinase1 gene (*Pink1-/-*) for testing at 3.5 and 9.5 months (mos) of age. The black asterisk shows that at 9.5 mos of age, rats in the *Pink1*-/- group made significantly more closed arm entries than WT rats; the white asterisk shows that *Pink1*-/- rats also made significantly more entries into closed arms than WT rats. (B) Bar graphs showing total amounts of time WT and *Pink1*-/- rats spent in the open arms (white), closed arms (black) and the center platform (striped) of the maze during testing at 3.5 and 9.5 mos of age. Asterisks show that at both 3.5 and 9.5 months of age, *Pink1*-/- rats spent significantly less time in closed arms and significantly more time on the center platform than WT rats. Stacked bar graphs showing percentages of total times *Pink1*-/- and WT rats spent on major behaviors within the open arms (C), closed arms (D) and center platform (E) during testing at 3.5 and 9.5 mos of age. Major behaviors examined included stationary behavior (black), ambulation (dark gray), rearing (light gray), grooming (white) head dipping (slanted stripes) and engaging in stretch-attend/scanning (vertical stripe) Significant group differences in the percentages of total times that rats devoted to a given behavior are marked by asterisks within the bar graphs of the group where significantly more time was spent.

At 9.5 months of age, a one-way ANOVA showed that WT rats made significantly fewer total arm entries compared to *Pink1*-/- subjects [5.71 vs. 8.87, F _(1,20)_ = 7.90, p= 0.011, □^2^ = 0.52, Fig 5A]. A repeated measures ANOVA further identified significant main effects of arm type [F_(1,20)_ = 145.21, p < 0.001, □^2^ = 0.88), a significant main effect of group [F_(1,20)_ = 8.70, p = 0.008, □^2^ = 0.30] and a significant interaction between these two [F_(1,20)_ = 5.95, p = 0.024, □^2^ = 0.23). These effects were driven by *Pink1*-/- rats entering closed arms nearly twice as often (7 vs. 5) as WT rats (p = 0.002, Fig 5A). In terms of times spent, a repeated measures ANOVA also revealed significant main effects of maze location [open arm, closed arm, center platform, F_(1.31, 26.20)_ = 109.26, p <0.001, □^2^ = 0.85] and a significant interaction between maze location and group [F_(1.31, 26.20)_ = 6.01, p = 0.015, □^2^ = 0.23). Follow-up pairwise comparisons showed that these effects were driven by WT rats spending significantly more time in closed arms (146 vs. 109 sec, p = 0.023) and significantly less time in the maze center (141 vs. 178 sec, p = 0.016) compared to the *Pink1*-/- group (Fig 5B). Rats of both genotypes spent roughly 12 sec in the open arms of the arena (Fig 5B). Finally, analyses of major behaviors exhibited in each portion of the maze found no significant main effects of behavior for open arms (□^2^ = 0.10). For the closed arms and center platform, significant main effects of behavior [F_(2.35-3.08, 47.01-61.53)_ = 12.61-217.62, p< 0.001 for both, □^2^ = 0.39-0.92] and significant interactions between behavior and group [F_(2.35-3.08,47.01-61.53)_ = 5.93-6.16, p < 0.001-0.003, □^2^ = 0.23-0.24] were found. For the closed arms (Fig. 5D), these effects were driven by WT rats spending roughly 9% less time ambulating (p = 0.008) and about 20% more time grooming compared to the *Pink1*-/- cohort (p< 0.001). For the center platform (Fig 5E), effects were driven by WT rats spending approximately 10% less time engaged in stretch attend/scanning (p = 0.008), about 5% less time rearing (p = 0.023) and 5-6% more time head dipping (p = 0.042) and ambulating (p < 0.001) compared to the *Pink1*-/- group.

### Grip Strength

Forelimb and hindlimb grip strength was measured in rats at 3.5 and 9.5 months of age. All measurements were normalized to total body weight. At both timepoints, the pull force exerted by forelimbs was greater than pushing/compressive force measured for hindlimbs in both rat groups (Fig 6A, B). However, there were no significant main effects of genotype/group on either of these measures at either age tested (forelimb: □^2^ = 0-0.11; hindlimb □^2^ = 0-0.03). There were, however, significant main effects of age on normalized grip strength measures for both groups [F_(1,23)_ = 8.21- 41.60, p <0.001- 0.009, □^2^ = 0.26-0.60]. These main effects were driven by increased normalized hindlimb grip strength forces in 9.5 compared to 3.5-month-old rats of both genotypes and increased forelimb grip strength for only WT controls.

**Figure 6.**
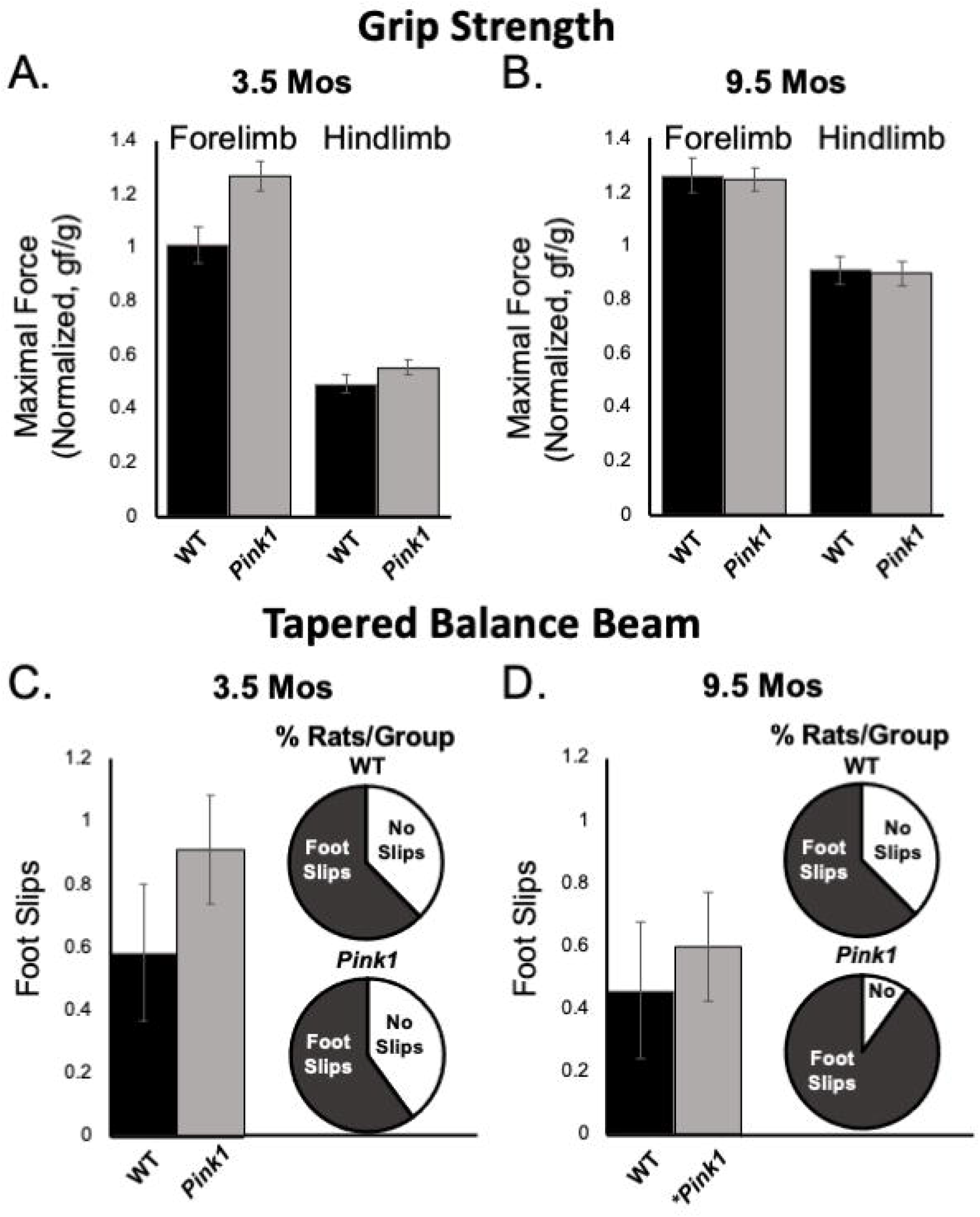
Grip Strength and Tapered Balance Beam data. Bar graphs showing measures of maximum forelimb (A) and hindlimb (B) grip strength/force normalized to body mass [grams of force (gf)/body weight in grams (g)] in wild type (WT, black bars) and rats with knockout of the PTEN (phosphatase and tensin homologue)-induced putative kinase1 gene (*Pink1-/-*, gray bars) measured at 3.5 and 9.5 months (mos) of age. There were no significant group differences in these measures of muscle strength at either age. Bar graphs showing average total numbers of foot slips committed by WT (black bars) or *Pink1*-/- rats (gray bars) in tapered balance beam traversal trials at 3.5 (C) and 9.5 (D) mos of age. Pie graph inserts show the proportion/percent of rats in each group that did or did not commit foot slips during the trial. Numbers of foot slips were minimal in both groups at both ages. However, proportionally more *Pink1*-/1 rats made foot slips compared to WT rats at 9.5 mos of age. The asterisk on the X-axis (D) denotes that five rats in the *Pink1*-/- group were removed from testing at 9.5 mos of age due to inability to cross the beam.

### Tapered Balance Beam

Tapered balance beam performance was assessed in rats at 3.5 (Fig 6C) and 9.5 months of age (Fig 6D). It is important to note that five of the *Pink1*-/- rats were no longer able to navigate the beam at the later time point and did not contribute to group data for this age. None of the WT rats were removed from these analyses. Average numbers of foot slips were slightly greater among *Pink1*-/- that did complete the task compared to WT rats at both ages. These differences, however, were small, not significant (□^2^ = 0.01-0.02) and represented small numbers of actual step offs/slips. The numbers/percentages of animals in each group that did and did not commit step offs/foot slips were also assessed; these percentages were similar among both groups of rats at 3.5 months of age (Fig 6C) but were greater in the Pink1-/- compared to WT rats at 9.5 months of age (Fig 6D).

## DISCUSSION

Cognitive impairments associated with PD are disabling for a considerable proportion of patients (Leroi, McDonald et al. 2012, Kudlicka, Clare et al. 2014, Rodriguez-Blazquez, Forjaz et al. 2015, Barone, Erro et al. 2017). These impairments are also resistant to most available treatments (Goldman and Weintraub 2015, Mack and Marsh 2017, Goldman, Vernaleo et al. 2018, Fang, Lv et al. 2020). Left unchecked, what may initially be mild deficits often progressively worsen and increase the likelihood of patients experiencing freezing and falls and developing PDD– which are leading causes of hospitalization, institutionalization and death among PD patients (Pigott, Rick et al. 2015, Peterson, King et al. 2016, Cholerton, Johnson et al. 2018). While *Pink1*-/- rats have been shown to recapitulate several key bio-behavioral aspects of PD, it has been largely unknown whether these rats also model the cognitive and/or memory sequelae associated with this disease. To address this question, the present study used three object recognition memory tasks to explore the face validity of this genetic rat model for the deficits in visual recognition memory and/or visuospatial information processing that commonly occur in PD patients (Owen, Beksinska et al. 1993, Higginson, Wheelock et al. 2005, Possin, Filoteo et al. 2008, Fang, Lv et al. 2020). These analyses revealed significant impairments in the *Pink1*-/- cohort in NOR, NOL and OiP performance as well as task-specific differences in the progression of these object discrimination/memory deficits across age. To summarize, at 3 months of age, rats of both genotypes showed robust ability to discriminate novel objects. The WT rats sustained these high levels of discrimination across all subsequent testing ages. However, in the *Pink1*-/- cohort, NOR performance declined sharply by 5 months of age and remained extremely low from this age on. In contrast, for the NOL task, rats of both genotypes initially (3 months of age) showed only modest ability to discriminate objects based on their location. However, by 5 months of age, NOL performance in WT rats rose to and remained at higher, more expected degrees discrimination while performance in the *Pink1*-/- group remained moderate to low at all ages. Finally, at 3 months of age the integrative recognition memory functions tapped in the OiP task were moderate in both WT and *Pink1-/-* rats. With successive testing, however, performance in WT rats incrementally increased and performance in *Pink1*-/- rats steadily declined. From these data it is tempting to speculate that knockout of the *Pink1* gene negatively impacts the brain circuits and/or neurochemical systems that are essential for performance in these tasks. However, given previous evidence for motor and affective disturbances in *Pink1*-/- rats (below) it is important to determine whether and how such non-mnemonic factors may have influenced the behavioral outcome measures observed. As discussed below, this was done by incorporating measures of motor function and affect/anxiety into analyses of object recognition task performance, and by bracketing the longitudinal object recognition testing sequence with elevated plus maze testing, measurements of hind- and forelimb grip strengths and assessments of motor coordination in traversing a tapered balance beam.

### Object Recognition in *Pink1*-/- Rats: Potential Confounds

As for many preclinical models of PD, it is important that analyses of object recognition memory testing take into consideration the possibility that motor and/or non-motor deficits could be confounding to data interpretation. For example, some muscular/motor effort is required for rats to get to and interact with the objects presented. In addition, disturbances in affect or anxiety can influence animals’ willingness to approach or explore objects, particularly those that are unfamiliar (Ennaceur, Michalikova et al. 2005, Ennaceur, Michalikova et al. 2006, Ennaceur, Michalikova et al. 2009). The previous studies showing that *Pink1*-/- rats experience progressive motor deficits and/or show an affective phenotype discussed below underscore the need for careful assessments to assure that the data from object recognition testing reported here reflect cognitive and/or mnemonic status. Accordingly, the present studies incorporated concurrent analyses of motor activity and affect into assessments of object recognition task performance, further evaluated motor function using a grip strength gauge and a tapered balance beam and further evaluated affect and anxiety using elevated plus maze testing.

#### Motor function

*Pink1*-/- rats are notable in part for progressive motor phenotypes. For example, this strain holds important and perhaps unique translational value for recapitulating early cranial and otolaryngeal sensorimotor deficits of PD(Kelm-Nelson, Lechner et al. 2021). As in PD patients (Ho, Iansek et al. 1998, Miller, Noble et al. 2006, Miller, Noble et al. 2006), *Pink1*-/- rats have been shown to have difficulty in sustained chewing and swallowing and show diminished vocalizing and vocalization volumes (Grant, Kelm-Nelson et al. 2015, Cullen, Grant et al. 2018, Kelm-Nelson, Brauer et al. 2018, Johnson, Kelm-Nelson et al. 2020). Other studies have shown that *Pink1* -/- rats also experience progressive somatic motor deficits. The most potentially concerning for the present studies are data identifying decreased novel open field locomotion and rearing, reduced hindlimb grip strength and increased commission of foot slips in traversing a tapered balance beam that in some (but not all) studies have been seen in *Pink1*-/- rats as young as 4 months of age (Dave, De Silva et al. 2014, Grant, Kelm-Nelson et al. 2015). In the present study, all rats were qualitatively evaluated for ability to freely locomote within the empty testing arena during the habituation/re-habituation trials that preceded every object recognition testing block. Although several *Pink1* -/- rats developed what appeared to be an uncoordinated gait at around 7-month-old, all were able to navigate the relatively small testing arena used and none were excluded from object recognition testing on this basis. Additional motor assessments made in conjunction with object recognition testing also showed that during test trials *Pink1*-/- rats often spent more time rearing and/or ambulating than WT rats. There were also no significant ‘before or after’ group differences in measures of fore- or hindlimb grip limb strength or commissions of foot slips on a tapered balance beam showed in *Pink1*-/- rats. It was noted, however, that *Pink1*-/- rats made slightly more foot slips than WT rats, that proportionally more *Pink1*-/- rats committed step offs than WT rats and that 5 of the *Pink1*-/- rats and none of the WT controls had to be removed from tapered balance beam testing at 9.5 months of age due to difficulty in remaining on the widest portions of the beam. Thus, we did find evidence of an emergent motor phenotype in the *Pink1-/-* cohort. However, further, more nuanced analyses are needed to resolve its nature. In the meantime, the qualitative and quantitative data in hand argue against motor impacts in the *Pink1*-/- group as interfering with object exploration or object recogntion testing. Importantly, the data also suggest that somatic motor deficits in *Pink1*-/- rats manifest later than do impairments in the cognitive and memory processes tapped in the NOR, NOL and OiP tasks. This could signal an additional dimension of face validity for the *Pink1*-/- rat model, as impairments in cognition and memory typically present during prodromal phases of illness, i.e., before the onset of measurable motor deficits, in PD patients (Caviness, Driver-Dunckley et al. 2007, Pigott, Rick et al. 2015, Aarsland, Creese et al. 2017, Baiano, Barone et al. 2020, Fang, Lv et al. 2020).

#### Anxiety/affect

Previous observations in *Pink1*-/- rats include behavioral measures suggesting increased anxiety (Kelm-Nelson, Trevino et al. 2018, Cai, Qiao et al. 2019, Hoffmeister, Kelm-Nelson et al. 2022). Such traits are potentially relevant for modeling aspects of mood disturbance that are common in PD—including clinical cases that are causally linked to loss of function *Pink1* gene mutations (Ephraty, Porat et al. 2007, Ricciardi, Petrucci et al. 2014). However, these traits could also adversely influence performance in object recognition testing. Specifically, while object recognition paradigms themselves are noted for provoking minimal stress or anxiety, baseline differences in anxiety can express as neophobia which reduces rats’ contact with objects–especially unfamiliar ones, and significantly erodes the discrimination indices typically used to quantify recognition memories (Ennaceur, Michalikova et al. 2006, Ennaceur 2010). Among the ‘other behaviors’ measured during object recognition testing were stationary behavior and grooming. The stationary behaviors observed were distinct from freezing. Accordingly, the significantly reduced times that *Pink1*-/- compared to WT rats spent stationary may be most likely to reflect diminished adaptation or habituation to the testing environment. The grooming that was observed occurred intermittently and included both cephalic and sequential grooming from head to body. Thus, interpretations with respect to decreased grooming in the *Pink1*-/- group leave it uncertain as to whether this difference reflects decreased or increased anxiety. To gain further clarity into this, rats were also tested on an elevated plus maze. Previous studies examining rats at 4 and 12 months of age showed that *Pink1*-/- rats entered and spent significantly more time in closed arms than controls (Hoffmeister, Kelm-Nelson et al. 2021). However, in the present study, *Pink1*-/- rats made more entries but spent less time in the closed arms than did WT rats. Further, while neither group spent much time in the open arms, *Pink1*-/- rats spent significantly more time in the center platform than WT rats. Finally, *Pink1*-/- rats spent significantly more time rearing and/or ambulating and less time grooming in the closed arms, and significantly more time engaged in rearing and stretch-attend/scanning and less time head dipping and ambulating in the center platform. Thus, the data are mixed with respect to behaviors classically aligned with increased or decreased anxiety. While these findings provide no indication of a *Pink1*-/- phenotype that would be likely to compromise object recognition testing, there is no question that there are significant differences in the ways in which *Pink1*-/- rats govern behaviors during object recognition and elevated plus maze testing compared to WT rats. Characterizing these differences more thoroughly and resolving their bases are important areas for future investigation.

### Impacts of Object Exploration in Time-limited Trials

The *Pink1*-/- rats assessed in this study were generated on a Long Evans background. Previous studies in this rat strain have demonstrated powerful effects of intermittent sample trial object exposure on subsequent discrimination of novelty. Specifically, it was shown that multiple, shorter exposures to sample objects greatly enhanced rats’ sensitivity to novelty demonstrated in test trials compared to a single, longer exposure period (Anderson, Jablonski et al. 2008, Shimoda, Ozawa et al. 2021). These findings drove the decision to incorporate multiple sample trials (3) in the testing protocols used here. Importantly, however, all trials were time-limited and thus subject to unintended impacts of differences in the time spent gaining familiarity with sample objects on later measures of memory strength or recall. Accordingly, analyses included evaluations of any group differences in total times rats spent with objects during both sample and test trial periods. These analyses showed that the generally more active state noted above in *Pink1* -/- compared to WT rats included knockout rats typically spending more to significantly more time actively exploring objects in all trial types. This argues against neophobia and argues against differential exposure to samples as negatively impacting measures of DI in the gene knockout group. The latter was further supported in findings of no significant or near significant positive correlations between the durations of sample or test trial object explorations and DI for any group for any task at any age. Careful analyses of sample trial object explorations also ruled out contributions of innate spatial bias or bias toward object type(s) as contributing to the group, task and age-specific patterns of differential object exploration/discrimination seen in test trials. Rather, as discussed further below, the data in hand may be explained by deleterious consequences of knockout of the *Pink1* gene for the brain circuits and neurochemical systems that mediate object recognition memory functions.

### Comparison to Previous Studies

To our knowledge, there has been only one previous assessment of cognition or memory in *Pink1*-/- rats. This study included Barnes maze and NOR testing as part of a larger *in vivo* brain imaging study that examined male rats at 6-8 months of age (Cai, Qiao et al. 2019). The data presented were in some cases limited. For example, because the data from Barnes maze testing were collapsed across trials, information about spatial working memory or spatial learning strategies was not available. However, measures of average daily latency to find the goal location showed no differences in performance within or across groups over four sequential testing days. Thus, rats of both genotypes appeared to learn and retain task information similarly. The latter is consistent with findings from other rodent models of PD that often do not recapitulate the long-term reference memory deficits that are characteristic of later stages of disease and PDD (Miyoshi, Wietzikoski et al. 2002, Da Cunha, Silva et al. 2006, Betancourt, Wachtel et al. 2016). For NOR testing, a single sample exposure (5 min) and a 60 min intertrial interval was used. While the *Pink1*-/- group showed no discrimination deficits, these data are difficult to interpret because–perhaps owing to the use of single sample trials, the control cohort showed no preference for novelty. Key methodological details were also lacking, including a description of habituation, information as to whether rats were tested during subjective days or nights, how object exploration was defined and measured and whether rats were tested before or after undergoing *in vivo* imaging. Thus, it is uncertain what may have driven the substantial differences between this prior and the present study where robust deficits in all object recognition memory domains assessed were present in *Pink1*-/- rats by 6-8 months of age.

The present studies used a longitudinal testing strategy to gain insights into the potentially progressive impacts of a PD-relevant gene perturbation on cognition and/or memory. This revealed diverging trajectories in object recognition memory testing performance in WT and *Pink1*-/- rats between 3 and 9 months of age. This was related in part to some unexpected evolutions in object recognition performance in WT rats across this span. Specifcally, for NOL testing, WT rats initially showed moderate levels of discrimination that jumped to much higher, asymptotic levels by 5 months of age. Similalry, for OiP, an initially moderate level of discrimination seen in testing at 3 months of age increased, albeit more incrementally, over the next 6 months. While developmental trajectories in object recognition memory performance have been noted, these are described for much younger rats and suggest that adult levels of performance are in place within the first month of life (Reger, Hovda et al. 2009, Ainge and Langston 2012, Westbrook, Brennan et al. 2014). Thus, the bases for the age-to-age differences noted in the WT rats of study are unclear. Importantly, however, the generally upward trajectory of their performances indicates that WT rats continued to engage in these tasks and were not negative affected by test-retest contingencies.

### Potential Substrates of Object Recognition Impairment in *Pink1*-/- Rats

Longitudinal testing showed that the *Pink1*-/- cohort examined developed robust discrimination deficits in NOR, NOL and OiP tasks according to task-specific timelines. These rats continue to be tested for motor function. Thus, direct pathophysiological correlates to these behavioral profiles are not available. However, previous multimodal *in vivo* magnetic resonance imaging (MRI) studies in *Pink1*-/- rats have identified significant changes in brain regions and circuits known to be critical for object recognition memories. For example, volumetric analyses have shown that areas including perirhinal and entorhinal cortex, dentate, subicular, CA1 and CA3 fields of the hippocampal formation, nucleus reuniens of the thalamus and several amygdaloid nuclei are significantly smaller in *Pink1*-/- compared to wild type rats (Cai, Qiao et al. 2019). Diffusion weighted MRI has also identified significantly decreased anisotropy in many of these same regions and resting state functional MRI has identified significantly reduced connectivity between neostriatum, midbrain DA regions, hypothalamus and thalamus and increased connectivity between ventral midbrain DA regions and hippocampus in *Pink1*-/- compared to wild type rats (Ferris, Morrison et al. 2018, Cai, Qiao et al. 2019). Together these findings show that many of the brain regions and networks known to be critical for object recognition memory (Aggleton and Nelson 2020, Barker and Warburton 2020, Barker and Warburton 2020, Chao, de Souza Silva et al. 2020) are vulnerable to the *Pink1* -/- genotype. In addition, although findings with respect to DA cell body loss have been variable (de Haas, Heltzel et al. 2019), NE cell loss, increased neostriatal concentrations of DA and decreased levels of basal and potassium-stimulated neostriatal release of DA, ACh and others have also been identified in *Pink1*-/- compared to control rats between the ages of 4 and 12 months (Dave, De Silva et al. 2014, Grant, Kelm-Nelson et al. 2015, Villeneuve, Purnell et al. 2016, Cullen, Grant et al. 2018, Creed, Menalled et al. 2019). Although little is currently known about the status of neurochemistry in other subcortical or cortical regions, these data nonetheless show patterns of dysregulation induced by the *Pink1*-/- genotype that involve neurotransmitters known to play pivotal roles in object recognition memories (Dere, Huston et al. 2007, Bus, Zizmare et al. 2020). Further, all of the indices of pathophysiology described above are present in *Pink1*-/- rats over time frames when the results of this study predict that significant impairments in multiple object recognition memory domains would be present. Future studies that combine *in vivo* imaging with behavioral analyses may be in an especially powerful position to map the progression of brain pathophysiology to the evolution of domain specific object recognition memory deficits. Although MRI analyses can be brain wide, current understanding of the points of overlap and divergence among the neural systems that underlie performance in discrete object recognition memory tasks can be used to generate and/or prioritize narrower, more specific hypotheses to be tested by these means.

## SUMMARY AND CONCLUSIONS

Novel object recognition, NOL and OiP testing continues to be extensively used to evaluate recognition memory and visuospatial information processing deficits that are similar to those experienced by PD patients in a range of different preclinical rodent models of disease (Grayson, Leger et al. 2015, Haghparast, Esmaeili-Mahani et al. 2018, Kyser, Dourson et al. 2019, Bharatiya, Bratzu et al. 2020, Boi, Pisanu et al. 2020, Kakoty, K et al. 2021). The present studies identified robust deficits in all three of these tasks in *Pink1*-/- rats. This is the first demonstration of face validity in this model for commonly occurring cognitive and memory impairments associated with PD. The longitudinal testing scheme used along with companion assessments of motor and affective function also showed that object recognition memory deficits in *Pink1*-/- rats progressively worsen and precede the onset of potentially confounding motor signs. The need for treatments that prevent or slow the course of cognitive or memory decline in PD-- and especially those that do so without interfering with treatment of motor signs, is urgent (Goldman and Weintraub 2015, Goldman, Vernaleo et al. 2018). The present findings of progressive cognitive and memory deficits along with the emergence of motor signs identify *Pink1*-/- rats as well suited for accelerating the pace discovery needed to fill this therapeutic gap. Key directions for future investigations using this model include assessments of potential face validity for the sex differences that are prominent in many aspects of PD including the incidence and severity of mild cognitive impairments (Janvin, Larsen et al. 2006, Cereda, Cilia et al. 2016, Liu, Locascio et al. 2017, Cholerton, Johnson et al. 2018, Oltra, Segura et al. 2021). The benefits of continued use of object recognition memory tasks for these purposes include their proven utility for evaluating sex and sex hormone impacts in rodent models of PD (Luine 2015, Costa, Sisalli et al. 2020, Lima, Meurer et al. 2021, Pinizotto 2022). This along with the undisputed value of these tasks in identifying candidate neural substrates (Dere, Huston et al. 2007, Brown, Barker et al. 2012, Aggleton and Nelson 2020, Barker and Warburton 2020, Chao, de Souza Silva et al. 2020) could ultimately help resolve points of common pathophysiological ground that render object recognition memories vulnerable not only in PD but also in other neurodegenerative disorders including Alzheimer’s disease and schizophrenia (Grayson, Leger et al. 2015).

## ACKNOWLEDGEMENTS

This work was supported by a Pilot Award from the Thomas Hartman Parkinson’s Research Center (to MFK) and by a grant from the National Institutes of Health (R21 NS11000, to MFK).

